# «Control of the phage defense mechanism by Quorum Sensing (QS) in clinical isolates of *Klebsiella pneumoniae*»

**DOI:** 10.1101/2023.12.05.570179

**Authors:** Antonio Barrio-Pujante, Inés Bleriot, Lucía Blasco, Laura Fernández-Garcia, Olga Pacios, Concha Ortiz-Cartagena, María López, Felipe Fernández Cuenca, Jesús Oteo-Iglesias, María Tomás

## Abstract

Multidrug resistant (MDR) bacteria and the shortage of new antibiotics are a serious health problem and have increased the interest in bacteriophages, with great potential as antimicrobial agents but they can induce resistance. The objective of the present study was to reduce the development of phage resistance in *K. pneumoniae* strains by inhibiting the Quorum Sensing (QS). The QS inhibition by cinnamaldehyde (CAD) was confirmed indirectly by the reduction of biofilm production and directly by a proteomic analysis. Also, the infection assays showed that the phage resistance mechanisms of the bacteria were inhibited when phage-resistant *K. pneumoniae* strains were treated with a combination of phages with CAD. Finally, these results were confirmed by proteomic analysis as proteins related to the phage defence such as CBASS (bacterial cyclic oligonucleotide-based anti-phage signalling) and R-M systems as well as tail fiber proteins were present under phage treatment but not with the combination.

**GRAPHICAL ABSTRACT:** 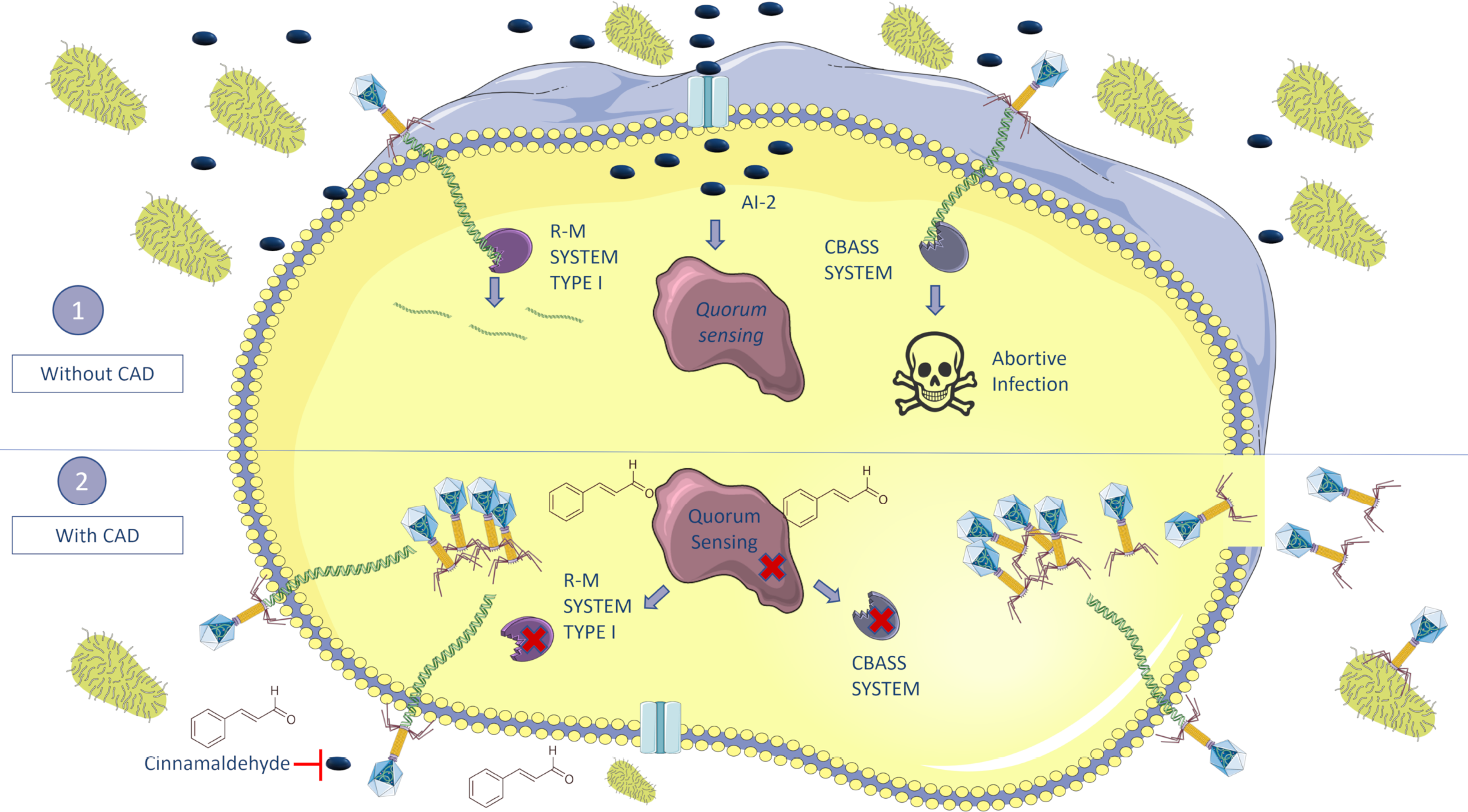

## INTRODUCTION

*Klebsiella pneumoniae* is a Gram-negative enterobacteria (1), catalogued by the World Health Organization (WHOs) in the list of ESKAPE pathogens (*Enterococcus faecium, Staphylococcus aureus, Klebsiella pneumoniae, Acinetobacter baumannii, Pseudomonas aeruginosa* and *Enterobacter)* that poses a serious threat to public health as multidrug resistant (MDR) bacteria and can be life-threatening for both, hospitalized patients and immunocompromised individuals. Moreover, many of them can persist stably through biofilm on catheters and ventilators as well as other medical devices (2).

In this context of clinical urgency, the use of phage therapy has recently re-emerged in the West as one of the main alternatives to treat MDR bacteria (3). Phages are defined as obligate intracellular parasites of bacteria (4). Phage therapy has several advantages over the use of antibiotic therapy: phages have high specificity of infectivity, infecting single bacterial species or subgroup of species, therefore are considered narrow-spectrum antimicrobials (3, 5); they don’t act on the patient’s normal microbiota; they are also highly effective against MDR pathogens and can be used in conjunction with antibiotics to restore its sensitivity, establishing synergies (6, 7); another advantage of phages is their ability to replicate inside the target cell at the site of infection. Finally, phages are easy to isolate and, being the most abundant biological entities on the planet, they can be an extraordinary way to obtain new low-cost antimicrobials (3, 5). However, the most important problem facing phages is the rapid acquisition of bacterial resistance, as they are in a constant “arms race” developing resistance mechanisms from both of them (3, 8).

The main mechanisms of bacterial phage resistance include: (i) outer membrane vesicles (OMVs), used as a decoy for phages to inject their DNA into the vesicule (9); (ii) blocking phage adsorption, avoiding the binding to the specific membrane receptor, i.e, by modification of this receptors through mutations (10) or masking them by capsule production or by biofilm formation (11); (iii) blocking DNA injection, i.e, superinfection exclusions (Sies system) (9); (iv) cut the injected DNA, if the phages still manage to enter the bacteria, there are defence mechanisms such as exogenous DNA cutting, like restriction-modification (R-M) (12) or CRISPR-Cas systems (13); (v) inhibiting phage DNA replication, i.e. bacteriophage exclusion system (BREX) or CBASS system (9, 14); (vi) interference with phage assembly, i.e, phage-inducible chromosomal island (PICI); (vii) abortive infection systems, in this resistance mechanism, bacteria when attempting to be infected by phage, causes its own death, thus protecting the rest of the bacterial population and preventing the spread of the infection, i.e. Toxin/Antitoxin systems (TA) (8, 9).

Many bacteria control the expression of defence mechanisms against phages through quorum sensing (QS) (15, 16), which is defined as a cellular communication process based on both production and secretion as well as detection of extracellular signaling molecules called autoinducers (AIs), which, depending on cell density, accumulate in the environment (17). QS allows to control of several processes such as fluorescence, virulence, biofilm formation, antibiotic resistance, bacterial competition factors, and phage resistance mechanisms like prophages, CBASS system, or CRISPR-Cas immunity (18-22).

The QS of *K. pneumoniae* relies mainly on the use of autoinducer-2 (AI-2), a furanosyl borate diester molecule encoded by LuxS synthase, for interspecies communication. Moreover, it is capable of detecting other autoinductors in the medium such as exogenous AHLs, known as autoinducers-1 (AI-1) (19, 21).

In this work, the role of QS in the control of the phage defence mechanisms was tested in *K. pneumoniae* in order to establish the basis of improvement of the phage therapy.

## MATERIAL AND METHODS

### Bacterial and phage strains

Two *K. pneumoniae* lytic phages were employed: the vB_KpnM_VAC36 phage (VAC36) (Family *Myoviridae*, Genus *Marfavirus*) and the vB_KpnM_VAC66 phage (VAC66) (Family *Myoviridae*, Genus *Slopekvirus*) (23). Both phage genomes, VAC36 (Genbank SAMN20298872) and VAC66 (Genbank SAMN22059211) are available from the GenBank Bioproject PRJNA739095.

3 clinical isolates of *K. pneumoniae* with different sensitivity to the phages tested were employed: K3318 resistant clinical isolate (GenBank SAMEA3649518), K3573 sensitive clinical isolate to phage VAC36 (GenBank SAMEA3649559), and ST974-OXA48 clinical isolate sensitive to phage VAC66 (GenBank WRWT00000000). All the clinical isolates of *K. pneumoniae* were clinical isolates from the Spanish hospital Virgen Macarena University Hospital (Seville, Spain) and the National Center for Microbiology (Carlos III Health Institute, Spain).

### Propagation and collection of phages

Phages used were propagated by the double-layer agar method (24). An overnight inoculum of *K. pneumoniae* isolation host of the phage was diluted 1/100 in LB medium and grew until OD_600nm_ 0.5. Later, 50 μL of phage were added to 200 μL of their *K. pneumoniae* isolation host and mixed with 4 mL of soft agar (0.5 % NaCl, 1 % tryptone, and 0.4 % agar) on TA agar plates (0. 5 % NaCl, 1 % tryptone and 1.5 % agar) and incubated at 37 °C during 24 h. TA agar plates are washed with SM buffer (0.1 M NaCl, 10 mM MgSO_4_, 20 mM Tris-HCl) and placed on a room temperature shaker for 3 h. Then, all liquid is recovered in 15 mL tubes and 1 % chloroform is added for 20 min. Finally, centrifuged for 15 min at 3400 x *g* and filtered through 0.45 nm filters.

### Cinnamaldehyde minimal inhibitory concentration assay

Cinnamaldehyde (CAD) (3-Phenylprop-2-enal; Sigma-Aldrich) minimum inhibitory concentration (MIC) for *K. pneumoniae* clinical strains K3318, K3573, and ST974-OXA48 was established by the chequerboard method (25). Briefly, in 96-well microtiter plates, nine serial double dilutions of CAD were prepared in Muller-Hinton broth (MHB). Each well was then inoculated with its corresponding *K. pneumoniae* strain to a final concentration of 5×10^5^ CFU/mL diluted from an overnight culture. A row of MHB inoculated with *K. penumoniae* was included as a positive control and a row including only MHB as a negative control. The plate was incubated for 24 h at 37 °C, and finally, the MIC was subsequently determined as the concentration of CAD in the first well where no bacterial growth was observed (25). All the experiments were performed in triplicates.

### Biofilm production

Biofilm production was used as an indirect measure of the QS and the QS inhibition was tested by the biofilm production in the presence of CAD (QS inhibitor). For this purpose, overnight inoculum of the clinical isolates of *K. pneumoniae* were diluted 1/10 (1×10^8^ CFU/mL) in modified Luria-Bertani (LB) medium (0.2 % Triptone; 0.1 % Yeast; 0.5 % NaCl) and 100 μl incubated in 96-well plates in presence of 3 different concentrations of CAD below the MIC (0.1 mM; 0.5 mM; 1 mM), for 24 h at 37 °C in darkness. A biofilm production without CAD was done as control. The next day, the biofilm production was quantified by the crystal violet staining method (26). Shortly, 100 μl of methanol were added to each well and was then discarded after 10 min. Once the methanol had completely evaporated, 100 μl of crystal violet (0.1 %) was added and the plates were incubated for 15 min. Finally, the wells were washed with PBS, and 150 μl of acetic acid (30 %) was added to resuspend the crystal violet adhered to the biofilm, and the absorbance was measured at OD_595nm_. All the experiments were performed in triplicates.

### Phage infection assays

Phage infection curves were developed with strains K3318, ST974-OXA48, and K3573, and phages VAC36 and VAC66.

Briefly, an overnight culture of the *K. pneumoniae* strain to be tested was diluted 1/100 in LB. Then it was allowed to grow to an OD_600nm_ of 0.3. Subsequently, four conditions were prepared in 96-well plates: growth control, *K. pneumoniae* culture in the presence of 1 mM CAD, *K. pneumoniae* strain in the presence of the corresponding phage at MOI 1 and, finally, a combination of the phage at MOI 1 and 1 mM CAD. The cultures were incubated at 37 °C by shaking for 24 h in a BioTek Epoch 2.0 (Agilent). The quantification of bacteria and phages was done by counts of colony-forming units (CFU/mL) and plaque-forming units (PFU/mL) at 0 hours (T0), 5 hours (T5) and 24 hours (T24).

To quantify the CFUs, 1 mL of culture was removed at the appropriate time and diluted to the correct dilution to count cells, and 100 μl of the dilution was plated on LB plates. For the quantification of PFUs, the phage must be isolated as follows. At the corresponding time points, 1 mL of culture was taken and 1 % chloroform was added, shaking it for 20 min. Then, it was centrifuged at 10000 x *g* for 5 min and serially diluted to the correct dilution to count. Finally, double-layer method was done to obtain of the PFUs (24).

The plates are incubated for 24 h at 37 °C and then CFU/mL and PFU/mL were quantified. All the experiments were performed in triplicates.

### Proteomic analysis

A proteomic study to determine the inhibition of phage defence mechanisms and QS was done by LC–MS and NanoUHPLC-Tims-QTOF analysis.

The K3318 strain (from an overnight inoculum) was inoculated into 50 mL flasks filled with LB medium at 1/100 dilution grown to an OD_600nm_ of 0.3 (around 10^7^ CFU/mL). The four conditions described previously were done. The culture strains were allowed to grow for 3 h. Next, 25 mL of each flask was removed to 50 mL tubes and placed for 10 min on ice and then centrifuged for 20 min at 4 °C and 4500 rpm, after discarding the supernatant, we freeze the pellet at -80 °C. The next day the pellet was resuspended in PBS medium and sonicated, then sonicated pellets were centrifuged for 20 min at 4 °C at 4300 x *g*. The supernatant was recovered and employed for the proteomic analysis. The quantitative analysis of proteins was done using 200 ng of supernatant for each sample. It was performed by NanoUHPLC-Tims-QTOF, the equipment used was a TimsTof Pro mass spectrophotometer (Bruker), a nanoESI source (CaptiveSpray), a QTOF-time analyzer and a nanoELUTE chromatograph (Bruker). Previous sample preparation was carried out by tryptic digestion in solution with reduction-alkylation followed by Ziptip desalting. Data were together in nanoESI positive ionization mode, Scan PASEF-MSMS mode and CID fragmentation mode, with an acquisition range of 100–1700 m/z. The products were separated on a Reprosil C18 column (150 × 0.075 mm, 1.9 μm, and 120 Å) (Bruker) at 50 °C, with an injection volume of 2 μL. The mobile phases consisted of 0.1 % H_2_O/formic acid (A) and 0.1 % acetonitrile/formic acid (B). The flow rate was 0.4 μL/min, and the gradient program was as follows: 11 % B (0–5 min), 16 % B (5–10 min), 35 % B (10–16 min), 95 % B (16–18 min) and 95 % B (18– 20 min). Finally, different software was used for data acquisition: Compass HyStar 5.1 (Bruker) and TimsControl (Bruker), DataAnalysis (Bruker), and PEAKS studio (Bioinformatics Solutions).

## RESULTS

### CAD MIC determination

The chequerboard assay was performed to determine the CAD MIC for clinical isolates of *K. pneumoniae*. The MIC (Figure 1A) for the clinical strain K3318 was 6.125 mM, for the clinical strain K3573 was 2.5 mM and for the clinical strain ST974-OXA48 was 3.125 mM.

**Figure 1.**
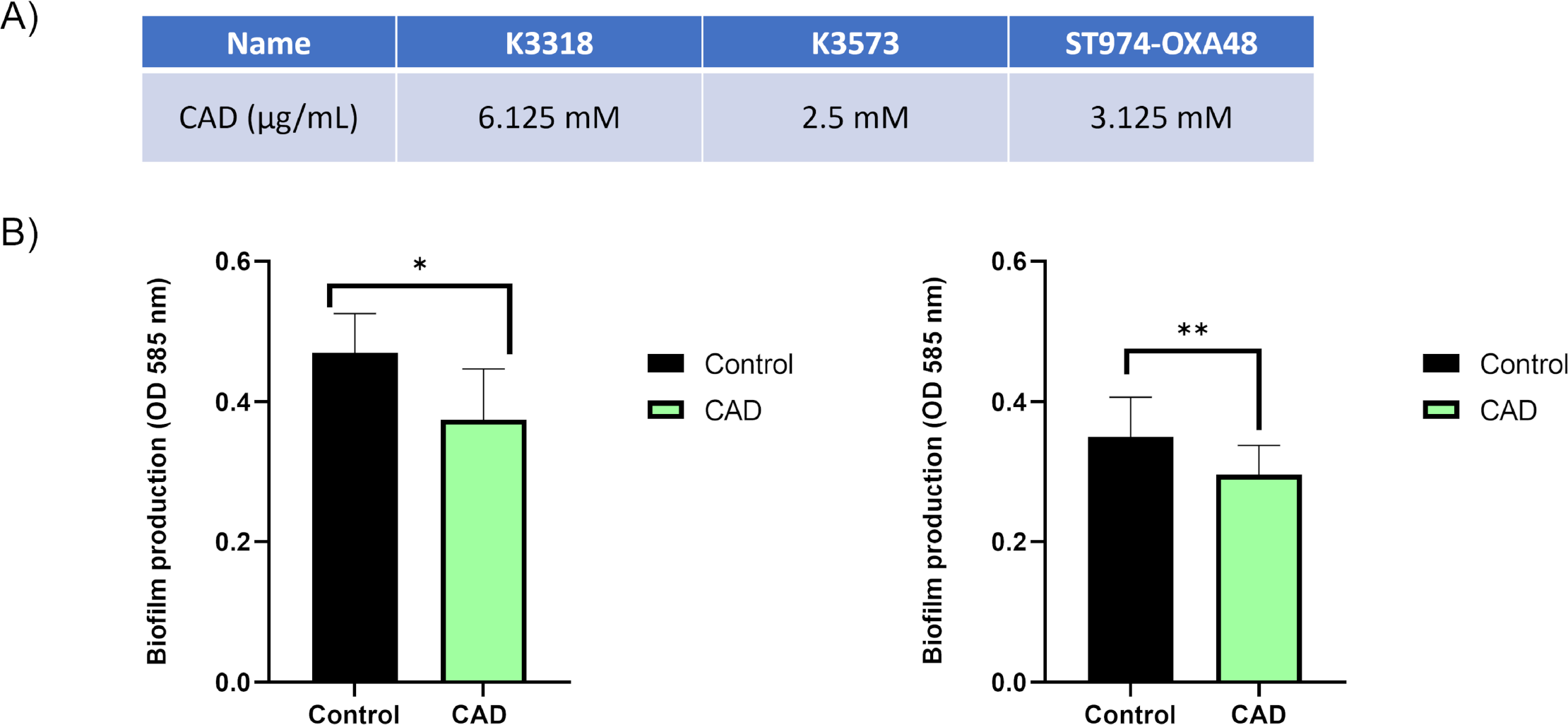
MIC for Cinnamaldehyde (CAD) and biofilm production in different clinical strains. A) MIC obtained for different strains of *K. pneumoniae* in presence of CAD. B) Significant decrease in biofilm production in the strains K3318 and K3573, in presence of 1mM of CAD respected to the control.

### Biofilm production assay

The biofilm production of *K. pneumoniae* clinical strains K3318, K3573, and ST974-OXA48 was tested as an indirect measurement of the QS activity (19, 27). The biofilm production was measured in the presence of different concentrations of the QS inhibitor CAD (0.1 mM, 0.5 mM, and 1 mM). The results showed a significant decrease in the ability to form biofilm in strains K3318 and K3573 in the presence of 1 mM CAD compared to the control (Fig. 1B), so this concentration was selected to inhibit the QS in the following assays. However, the strain ST974-OXA48 did not produce biofilm (OD_585nm_, 0.05).

### Phage infection assays

Phage infection curves were performed to precise the infectivity of phages in the presence of the QS inhibitor CAD to determine if the phage defence mechanisms are controlled by QS. Two phages were employed, VAC36 and VAC66, and 3 clinical isolates of *K. pneumoniae*, the clinical isolate K3318, resistant to both phages and the clinical isolates K3573 and ST974-OXA48, sensitives to VAC36 and VAC66 respectively.

The infection curves showed a decrease in the optical density of the strain K3318 when the infection was done with the phages in combination with CAD (Fig. 2A and B), as this strain was resistant to both phages, in the infection control curve the growth was similar to the control. In the case of the sensitive strains, no differences were observed when the infection was done with the phage alone or in combination with CAD (Fig. 2C and D). To confirm the results observed, counts of CFU and PFU were done at 0 h, 7 h, and 24 h (Fig. 3A and B). The results of these counts showed a significant decrease at 7 h in the number of CFUs and a significant increase in the PFUs counts for strain K3318 when infected with the combination of CAD and each of the phages, this reduction of CFUs and increase in PFUs was not observed when the infection was done with each phage alone. At 24 h the number of CFUs was increased in this condition but it was significantly lower than the control with phage and the growth control. In the same way, at 24 h the PFUs quantified were diminished but they were significantly higher than those quantified for each phage alone. These results are consistent with a productive phage infection when the CAD was present, suggesting that CAD is favouring the infection by inhibiting the phage defence mechanisms.

**Figure 2.**
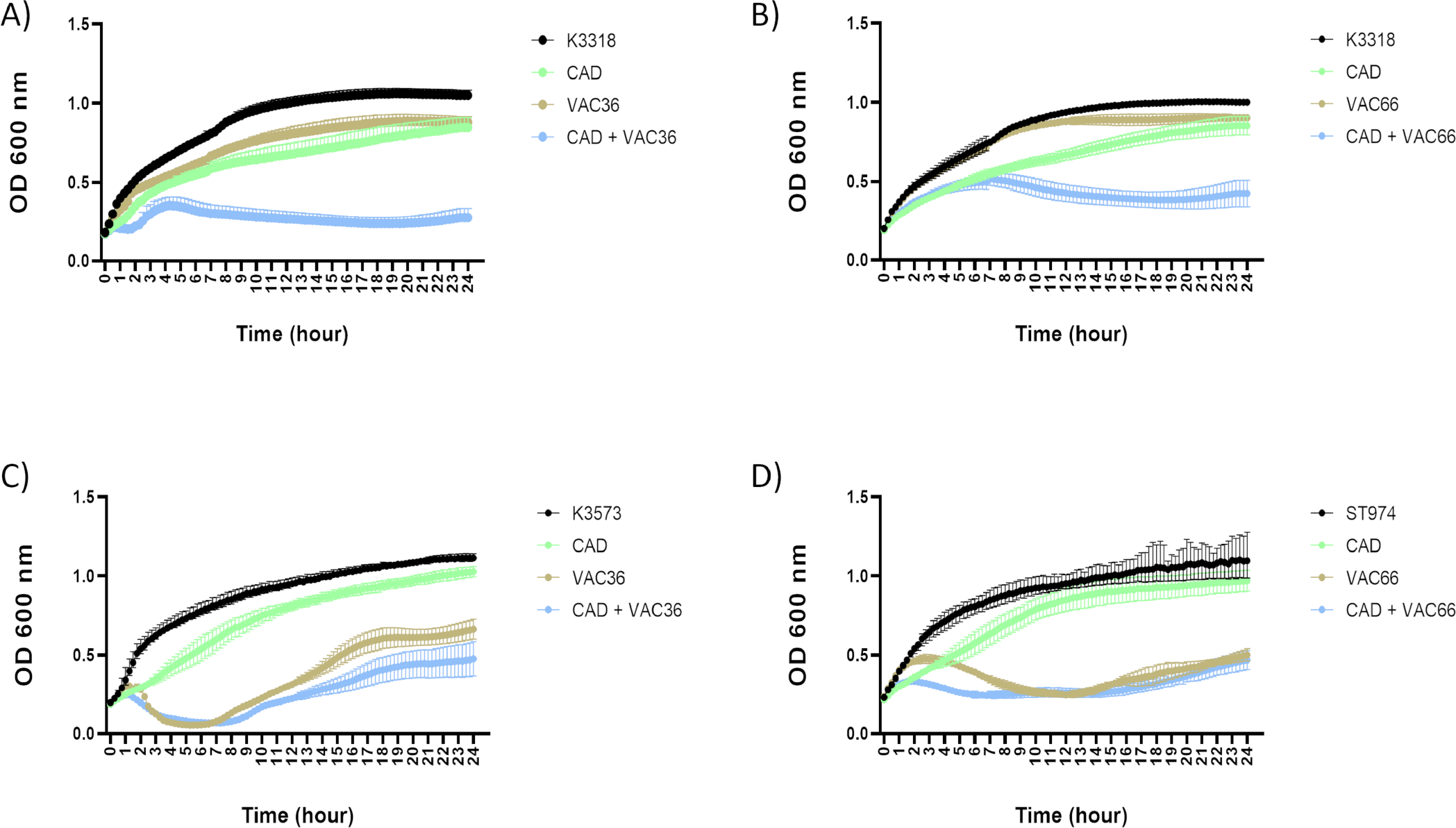
Infection curves with the three clinical isolates of *K. pneumoniae* with phage alone and in combination with CAD. A) Strain K3318 infected with VAC36. B) Strain K3318 infected with VAC66. C) Strain K3573 infected with VAC36. D) Strain ST974-OXA48 infected with phage VAC66.

**Figure 3.**
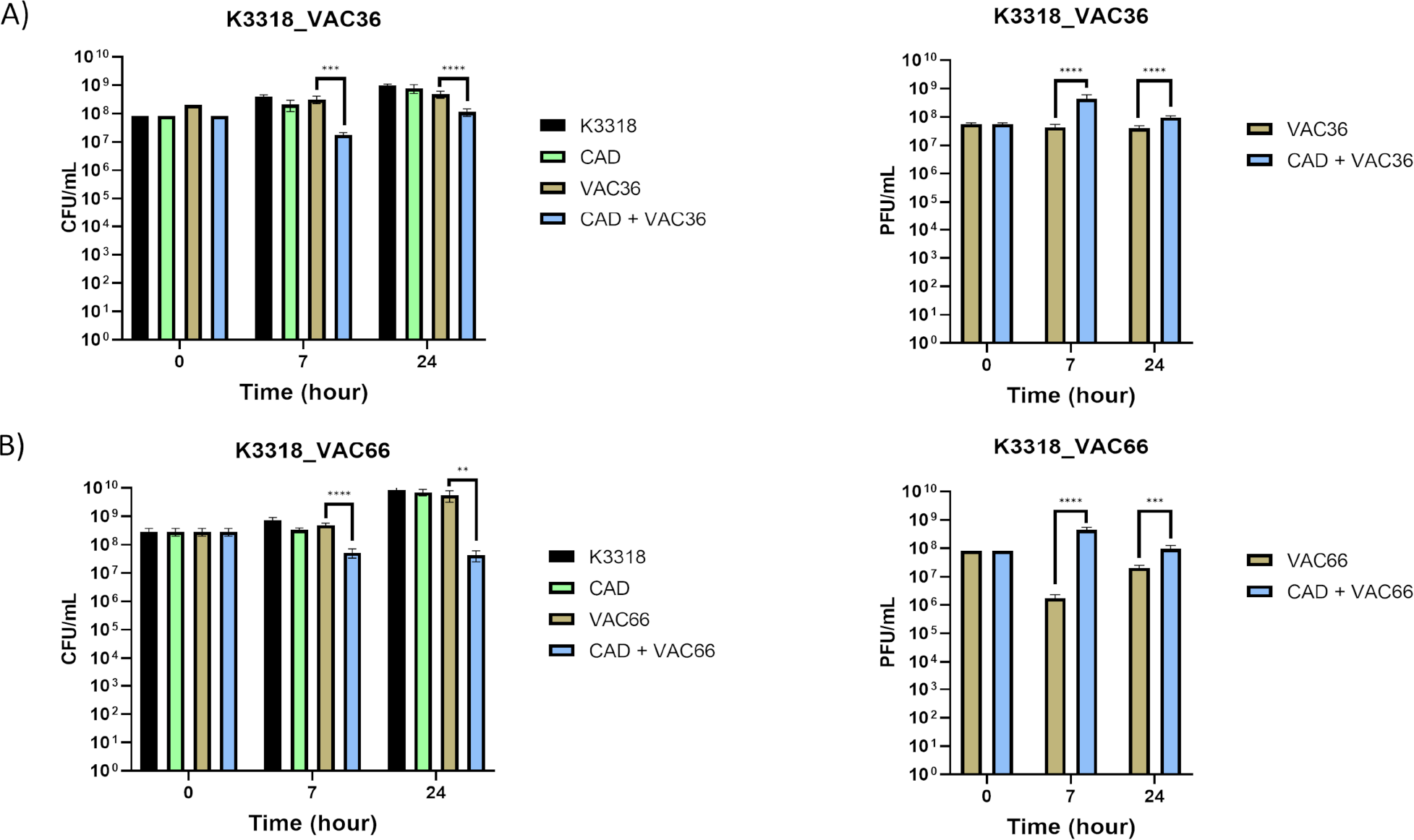
Quantification of bacteria and phages by CFU/mL and PFU/mL counts in phage infection assays. A) CFU/mL and PFU/mL of strain K3318 infected with phage VAC36. B) CFU/mL and PFU/mL of strain K3318 infected with phage VAC66.

Host strains K3573 and ST974-OXA48 were sensible to phage infection but no significant differences were found between treatment with phage alone and with the combination (Fig. 1C and D). As no differences between phage and phage combined with CAD were observed, CFUs and PFU were not quantified. These results showed that in these strains no defence mechanisms were active under any of the infection conditions, as infection occurs always in the presence of phage.

### Study of the proteins related to phage defence mechanisms and QS

To confirm that the phage defence mechanisms are controlled by the QS, a proteomic analysis by NanoUHPLC-Tims-QTOF was done. The analysis identified 231 proteins from the CAD combined with VAC36 condition and 169 proteins from the condition with phage alone, corresponding to 15 functions (Fig. 4A and B).

**Figure 4.**
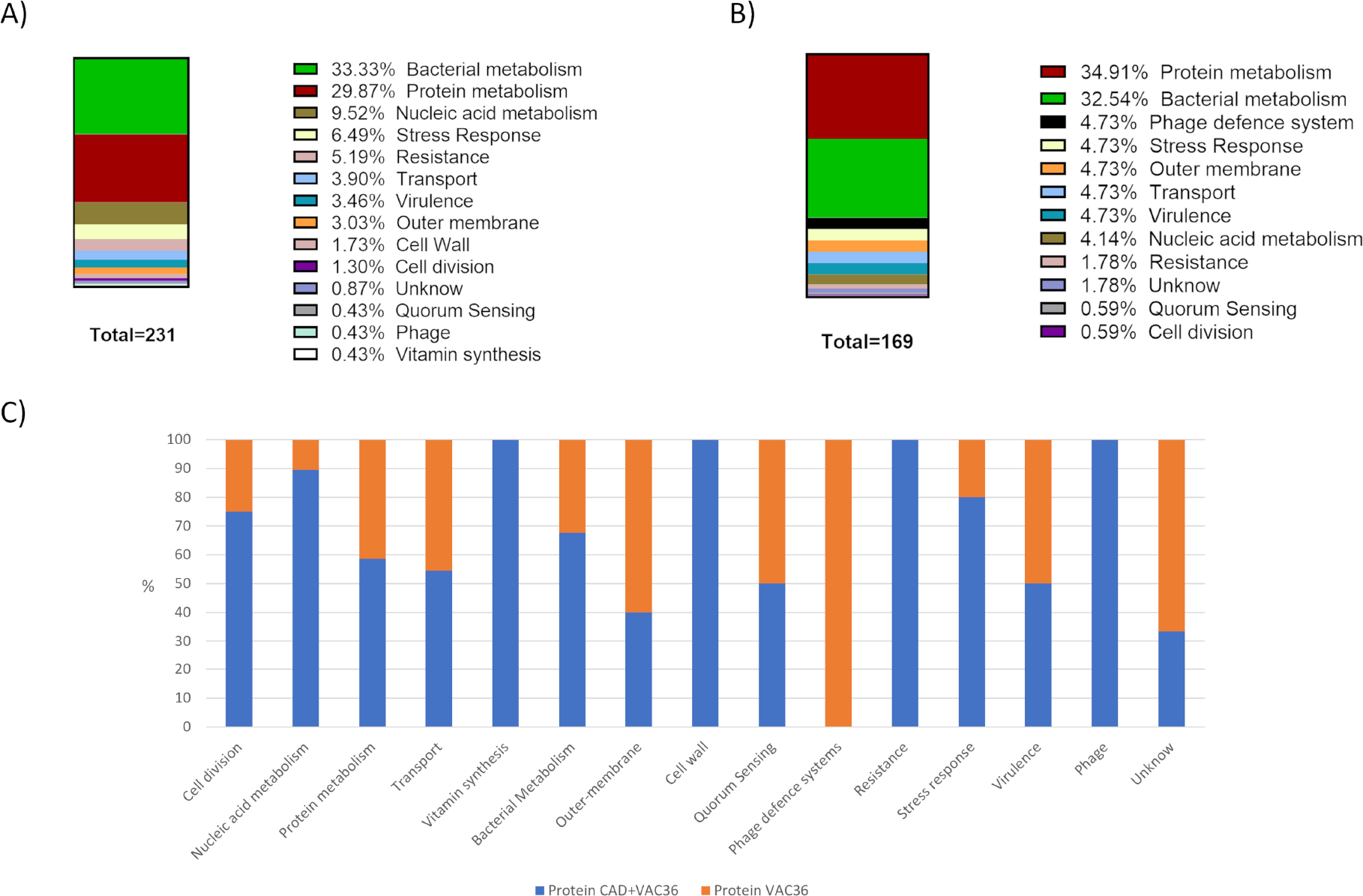
Graphical representation of proteomic analysis. A) Abundance of proteins belonging to each functional group in the CAD combined with VAC36 condition. B) Abundance of proteins belonging to each functional group in the VAC36 condition. C) Relative abundance of each functional protein group in K3318 infected with phage VAC36 and with phage VAC36 in combination with CAD. Parts of the figure were drawn by using pictures from Servier Medical Art. Servier Medical Art by Servier is licensed under a Creative Commons Attribution 3.0 Unported License (https://creativecommons.org/licenses/by/3.0/)

The analysis showed that the main representative functions in both conditions were those related to bacterial metabolism and to protein metabolism (Fig. 4A and B). The relative abundance of functions related to acid nucleic metabolism was slightly higher on CAD combined with VAC36 condition (Fig. 4C). Interestingly, the analysis showed a higher relative abundance of proteins related to the phage defence when the CAD was absent. QS-related proteins were found to have a lower relative abundance in the presence of CAD, confirming the QS inhibitory role of this compound (Table 1).

**Table 1.** Proteins found in the proteomic analysis associated with Quorum Sensing (QS).

| QUORUM SENSING |  |  |  |  |  |  |
| --- | --- | --- | --- | --- | --- | --- |
| Description | Accession no. | -10LgP | Control | CAD | Mechanism | Ref. |
| methionine adenosyltransferase | UZL31645.1 | 318.0766 | 1.51E+03 | 3.89E+02 | AI-2 Pathway | (47) |
| autoinducer 2 ABC transporter<br>substrate-binding protein LsrB | UZL28532.1 | 236.32896 | 8.80E+02 | 2.24E+02 | AI-2 Pathway | (17) |

As shown in Table 2, the proteins related to phage defence mechanisms were highly expressed when the infection was done with the phage alone but not when the infection was done in the presence of CAD, thus confirming the relation of QS and phage defence mechanisms. In addition, a tail phage protein was present only when the infection was done in the presence of CAD, confirming the active infection.

**Table 2.** Proteins found in the proteomic analysis associated with phage defence mechanisms and phage proliferation.

| PHAGE DEFENCE |  |  |  |  |  |  |
| --- | --- | --- | --- | --- | --- | --- |
| Description | Accession no. | -10LgP | CAD+VAC | VAC | Mechanism | Ref. |
| purine-nucleoside phosphorylase | UZL32988.1 | 310.17953 | 0.00E+00 | 3.09E+03 | CBASS System | (14) |
| uridine phosphorylase | WGN84707.1 | 187.73683 | 0.00E+00 | 2.42E+02 | CBASS System | (14) |
| restriction endonuclease subunit S | UZI85743.1 | 60.279945 | 0.00E+00 | 2.32E+03 | R-M System | (42) |
| ubiquitin-like protein, partial | WP_289465582.1 | 132.11082 | 0.00E+00 | 6.72E+02 | CBASS System | (39) |
| polyubiquitin, partial | WP_223807730.1 | 201.96265 | 0.00E+00 | 6.72E+02 | CBASS System | (39) |
| PHAGE PROTEIN |  |  |  |  |  |  |
| Description | Accession no. | -10LgP | CAD+VAC | VAC | Mechanism | Ref. |
| tail fiber domain-containing protein | WP_153932364.1 | 172.86615 | 4.65E+02 | 0.00E+00 | Tail phage | (43) |

## DISCUSSION

Phages are a great opportunity for the future in the war against MDR bacteria, they can be used alone, in cocktails, or in combination with antibiotics as they can restore sensitivity to them (28). Phages have many advantages over the use of conventional antimicrobials, they are easy and cheap to obtain, they have high specificity, as well as being self-replicating (3, 5-7). In this context, the present study aimed to determine if phage resistance mechanisms are controlled by QS. To achieve this objective, several phage infections were conducted in the presence of the CAD, a QS inhibitor. Previous studies have shown that QS controls phage defence mechanisms in bacteria because under conditions of high cell density, they are more exposed to phage attack as phages are more abundant in densely populated environments, as they necessarily need bacteria to proliferate (29).

In this work, the QS inhibition in clinical isolates of *K. pneumoniae* was confirmed by the reduction of biofilm when the CAD was present, except for a non-producing biofilm strain, the ST974-OXA48. As CAD was confirmed as a QS inhibitory compound for *K. pneumoniae*, the role of QS in the expression of phage defence mechanisms was tested by infecting the phage-resistant strain *K. pneumoniae* K3318 with two different phages in combination with CAD 1mM. The infection curves performed showed a significant reduction in OD_600nm_ when resistant strain K3318 was infected with both phages in the presence of CAD, as well as a reduction in CFU/mL and PFU/mL at 7 h and 24 h, the infection controls were very similar to the growth control values. In the sensitive strains, no defence mechanisms were activated so the infection was similar when CAD was present and absent. These results suggested that when QS was inhibited the defence mechanisms were also inhibited. Several previous works confirm these results, in 2016, Hoque MM *et al*. demonstrated that *V. cholerae* QS, which uses cholera AI-1 and AI-2-like autoinducers, controls both the production of haemagglutinin protease (HAP) which was responsible for inactivating viral particles and the deregulation of phage receptors, in particular the LPS O-antigen receptor, preventing phage adhesion (30). Moreover, in 2023 Severin GB *et al*., associated the high cell density in *V. cholerae* cultures with the transcription of two essential components of the CBASS defence system, the oligonucleotide cyclase and the effector phospholipase (20). Regarding the CRISPR-Cas defence system, numerous studies confirmed their activation at high cell density, so in *Serratia* spp. there was an increase of type I-E, I-F, and III CRISPR-Cas systems (31), and also in *P. aeruginosa*, the QS was demonstrated to be involved in the regulation of CRISPR-Cas genes and the activation of three key aspects of the system, expression, activity, and adaptation (15, 22). In addition, in 2021 Shah M *et al*., suggested that *P. aeruginosa* controls the cell death by QS quinolone signal (PQS) to prevent the spread of phage infection (32). In *Escherichia coli*, it was demonstrated that the activation of QS reduced the number of phage λ receptors on the cell surface in order to evade the infection (29). Finally, the BREX defence system was also ligated to the QS control as the S-Adenosyl Methionine (SAM), which is a precursor in the synthesis of AI-2, could be a necessary cofactor for the BREX system (17, 33). To our knowledge, we could not find studies specifically linking *K. pneumoniae* QS to phage defence mechanisms such as those previously described.

In this work, the relation between the QS and phage defence mechanism was tested by the inhibition of QS. CAD was selected as a QS inhibitory compound, as it was previously described to interfere with the AI-2 based QS *Vibrio spp*. and *Burkholderia spp*. by decreasing the DNA binding activity of the response regulator LuxR (34-36). As for *Vibrio spp*. the QS autoinductor in *K. pneumoniae* is the AI-2 synthesized by an ortholog of LuxS synthase and related with the production of biofilm (37), a relation also observed in *E. coli* uropathogenic strains in a study in which the use of CAD reduced the QS and the biofilm production (38).

The results of the proteomic study confirmed both the results obtained in the infection curve and the relation between QS and phage defence mechanisms. The relative abundance of proteins related to DNA functions was a little higher when the infection of VAC36 was done in the presence of CAD (Fig. 4C), probably because the assay was done at 3 h, and at this point, the bacteria was in an exponential growth phase (Fig. 2A). It was also observed the presence of a tail phage protein when the QS was inhibited, suggesting the synthesis of new virions that together with the absence of phage defence related protein is indicative of the occurrence of an active infection (Table 2). When the infection was done with the phage but without inhibiting the QS, no phage proteins were observed and several defence proteins were present (Table 2), which corresponded with the absence of infection observed in the infection curves (Fig. 2A). Many of the proteins related to phage resistance belonged to the CBASS defence system, an abortive infection system that acts first through an oligonucleotide cyclase, activated when the phage infects the bacterium, and a cyclic oligonucleotide-sensitive effector that kills the infected cell through its activity (14, 20, 39). The CBASS proteins identified were purine-nucleoside phosphorylase, uridine phosphorylase, ubiquitin-like protein, and polyubiquitin, the two first proteins act in the CBASS as effector proteins as both are purine nucleoside phosphorylases (PNP) (14), and the ubiquitin and polyubiquitin act as part of the coding of Cap2, a gene whose catalytic activity is essential for the correct functioning of the CBASS system (39). Another protein that was absent in the presence of CAD and related to the restriction-modification system (R-M) defence system was the restriction endonuclease subunit S (40). The R-M is a type of innate immunity of prokaryotes that protects cells from the insertion of foreign DNA, that includes a restriction endonuclease (Rease), which recognizes short DNA sequences, and cuts them, and a methyltransferase (Mtase), which methylates host DNA so that it cannot be cut by Rease (41). The restriction endonuclease subunit S belongs to the R-M type I system, which is encoded by three genes: hsdR encoding the restriction subunit (R), hsdM the modification subunit (M), and hsdS the recognition subunit (S for specificity) (42). Furthermore, a phage tail protein was found only when phage and CAD were combined, which assures that a productive infection was taking place in this condition and new phage progeny was being synthetizing (43).

Moreover, the QS inhibition with the CAD was validated by the lower expression of some QS proteins when CAD was added in comparison with the control. Some of these proteins were directly related to the synthesis or regulation of the AI-2 autoinducer production, methionine adenosyltransferase (MAT), a key bacterial enzyme involved in the cascade of regulation and formation of AI-2, the QS autoinducer of *K. pneumoniae* (17, 44); and autoinducer 2 ABC transporter substrate-binding protein LsrB, responsible for transporting AI-2 into cells (17, 45). Finally, in the CAD and phage condition, the activation of multiple resistance mechanisms like efflux pumps was observed (Fig. 4C), this is probably a consequence of the presence of CAD, in *P. aeruginosa*, Tetard A. *et al*., showed that exposure to sub-inhibitory concentrations of CAD resulted in the expression of efflux pump encoding operons (46).

In conclusion, as the inhibition of QS significantly reduces phage resistance, its therapeutic use could be considered in combination with phage cocktails or introducing the QS inhibitor compound in the phage-antibiotic combination, which could further enhance the synergy and reduce the appearance of resistance, solving part of the deficiencies that phages have on their own. For all this, the inhibition of QS can be the piece of the puzzle that completes the great potential of phages, and in turn we can find the solution to a problem that is increasingly approaching, MDR bacteria and the lack of antibiotics.

## FUNDING

This study has been funded by Instituto de Salud Carlos III (ISCIII) through the projects PI19/00878 and PI22/00323 and co-funded by the European Union, and by the Study Group on Mechanisms of Action and Resistance to Antimicrobials, GEMARA (SEIMC). (SEIMC, http://www.seimc.org/). This research was also supported by CIBERINFEC (CIBER21/13/00095) and by *Personalized and precision medicine* grant from the Instituto de Salud Carlos III (MePRAM Project, PMP22/00092). M. Tomás was financially supported by the Miguel Servet Research Programme (SERGAS and ISCIII). O. Pacios, L. Fernández-García and M. López were financially supported by the grants IN606A-2020/035, IN606B-2021/013 and IN606C-2022/002, respectively (GAIN, Xunta de Galicia). I.Bleriot was financially supported by the pFIS program (ISCIII, FI20/00302). Finally, to thank to PIRASOA laboratory which is the reference laboratory for molecular typing of nosocomial pathogens and detection of mechanisms of resistance to antimicrobials of health interest in Andalusia, Virgen Macarena Hospital, Seville, to send us the clinical isolates. Thanks to Alvaro Pascual and Luis Martínez-Martínez from Virgen Macarena Hospital, Seville and Reina Sofia Hospital, Cordoba.

## AUTHOR CONTRIBUTIONS

A.B-P. and I.B., conducted the experiments and write the manuscript; L.B., L.F-G. and O.P., supervised the experiments; C.O-C. and M.L., helped to development the experiments. F.F-C. and J.O-I., collaborated in the edition of the work, and finally, M.T., supervised the experiments, validated of the results and financed the work.

## TRANSPARENCY DECLARATIONS

The authors declare not to have conflict of interest.

## Notes

### Competing Interest Statement

The authors have declared no competing interest.

